# Chemosensory afference in the tentacle nerve of *Lymnaea stagnalis*

**DOI:** 10.1101/2022.11.28.518212

**Authors:** Carmen C. Ucciferri, Russell C. Wyeth

**Affiliations:** Department of Immunology, University of Toronto, Toronto, Ontario, M5S 1A8, Canada; Department of Biology, St. Francis Xavier University, Antigonish, Nova Scotia, B2G 2W5, Canada

**Keywords:** *Lymnaea stagnalis*, neuroethology, chemosensation, gastropod, odours

## Abstract

Although the neural control of behavior has been extensively studied in gastropods, basic gaps remain in our understanding of how sensory stimuli are processed. In particular, there is only patchy evidence regarding the functional roles of sense organs and the extensive peripheral nervous system they contain. Our goal was to use extracellular electrophysiological recordings to confirm the chemosensory role of the tentacles in the great pond snail, *Lymnaea stagnali*s. Employing a special twin channel suction electrode to improve signal-to-noise ratio, we applied three food odours (derived from earthworm-based food pellets, algae-based pellets, and fresh lettuce) to a reduced preparation of the tentacle while recording neuronal activity in the tentacle nerve. Responses were assessed by comparing average spike frequencies produced in response to saline flow with and without odours. Confirming the omnivorous nature of this gastropod, we report strong neuronal responses to earthworm-based food odours and mild neuronal responses to algae-based food odours. There were no clear neuronal responses produced when lettuce food odour or control saline was applied to the tentacle. Overall, our results provide strong evidence for the chemosensory role of the tentacles in *L. stagnalis*. While it is unclear whether the differences in neuronal responses can be explained by differing sizes, numbers, or populations of neurons, these results are a useful foundation for further study of peripheral nervous system function in gastropods.

**Summary Statement:** Great pond snail tentacles send sensory signals to the brain in response to some (but not all) food odours.

## Introduction

Gastropods have been widely used for studying the neural control of behaviour. Components of sensory systems, central processing, and motor control have all been studied in a range of different species (Chase 2002). However, a major gap in this work has been in fully understanding the peripheral components of the sensory systems (Croll 2003). Numerous studies have explored the neuroanatomy and functional roles of gastropod cephalic sensory organs (e.g., Fredman and Jahan-Parwar 1980; Chase and Tolloczko 1993; Gobbeler and Klussmann-Kolb 2007; Wollesen et al. 2007; Wyeth and Croll 2011; McCullagh et al. 2014). Yet, the sensory modalities remain unknown for the abundant and varied putative peripheral sensory cell types, as does much of the peripheral neural circuitry and functional integration with the central nervous system (Wyeth 2019). *Lymnaea stagnalis* is a particularly well studied gastropod, both with regard to its neuroethology, as well as many other aspects of its biology (Benjamin 2008; Koene 2010; Getz et al. 2018; Fodor et al. 2020; Kuroda and Abe 2020; Rivi et al. 2020; Rivi et al. 2021). Even in *L. stagnalis*, though, there are basic gaps in knowledge regarding the peripheral nervous system. Although chemosensation is presumed to be a primary function of gastropod cephalic sense organs, this has been experimentally verified in relatively few species (Wyeth 2019), and there is no direct evidence for a chemosensory role for the tentacles in *L. stagnalis* (although see Bovberg 1968). Thus, the goal of our study was to use electrophysiological techniques to record chemosensory responses to various food odours in the tentacle nerves of *L. stagnalis*.

*L. stagnalis* is an aquatic pulmonate gastropod that can be found across Europe, North America, and Asia in both slow-moving and stagnant freshwater bodies of waters (Kuroda and Abe 2020). These gastropods are omnivorous, feeding primarily on plants and algae, but will feed on carrion if available (Bovberg 1968). Foraging in most gastropods involves odour-based navigation, potentially involving one or more distinct strategies: kinesis (random movements modulated by odour concentrations changes), chemotaxis (following chemical gradients), and odour gated rheotaxis (following the flow when odours are present) (Croll 1983; Wyeth 2019). A recent study investigating the navigational behavior of *L. stagnalis* to attractive odours suggests that either chemotaxis or odour-gated rheotaxis are the strategies used to locate food (Alansari 2021). In either case, successful navigation in response to odour cues requires chemosensation. Given their anterior position above the substrate, the tentacles are an obvious candidate sense organ used to detect the food odours used during navigation towards distant odour sources. Several lines of evidence further support the expectation that the tentacles will provide chemosensory afference during navigation towards odour sources. Neuroanatomical studies of gastropod cephalic sense organs have revealed an extensive array of putative sensory cells (e.g., Emery and Audesirk 1978; Xin et al. 1995; Nakamura et al. 1999; Gobbeler and Klussmann-Kolb 2007; Wyeth and Croll 2011; Horvath 2020). In *L. stagnalis*, immunohistochemical staining and *in situ* hybridization revealed the presence of peripheral neurons distributed across both the ventral and dorsal edges of the tentacles and lips (Wyeth and Croll 2011; Young et al. 2022). Although modalities of the cell types are not known, one or more of them are presumed to be chemosensory. In the nudibranch *Tritonia exsulans* (previously *T. diomedea*), lesioning the rhinophore (anterior cephalic sense organs homologous to the tentacles in *L. stagnalis)* or their nerves eliminated the ability to navigate toward attractive odour sources (Wyeth and Willows 2006). Thus, there is reasonable evidence suggesting the tentacles will contain chemosensory cells that will respond to food odours in *L. stagnalis*.

Despite the accumulated evidence, studies to date have not found any chemosensory afference in the tentacle nerve of *L. stagnalis*. Rather, efforts have exclusively focused on responses to feeding stimulants, particularly sucrose, which only appears to produce afference in one of the lip nerves (Goldschmeding and Jager 1973; Kemenes et al. 1986; Kemenes 1994; Nakamura et al. 1999). It is certainly possible that while feeding stimuli act through chemosensory cells in the lip, other chemical stimuli originating from distant odour sources could be detected by the tentacles. Moreover, electrophysiological studies of the cephalic nerves of gastropods are challenging, given the small signals that will be produced from the small diameter axons associated with the putative peripheral sensory cells. Therefore, we hypothesized that the tentacle nerves would indeed carry chemosensory afference in response to the odorants known to produce navigational behaviours. Verifying this functional role for the peripheral nervous system in the tentacles is an initial step towards both characterizing the modalities of the different putative sensory cells and also determining the preliminary processing of sensory responses that may occur in the periphery. Thus, our aim was to use a modified *en passant* electrode to improve the signal-to-noise performance of recordings of neuronal activity in the tentacle nerve of *L. stagnalis*, and thereby test for chemosensory afference in response to three different odour sources known to produce differing navigational responses in *L. stagnalis*.

## Methods

### Animals

All animal use was consistent with Canadian Council for Animal Care guidelines and approved by the animal care committee at Saint Francis Xavier University. *L. stagnalis* were housed in freshwater tanks kept at room temperature and a photoperiod set to match the ambient outdoor cycle. The snails were fed a diet of Romaine lettuce and spirulina fish food daily, with commercially available fish pellet food given once weekly. Chalk was also added to the tanks to support stronger shell growth.

### Dissection

Snails were anaesthetized in a well-mixed solution of 125 *μ*L 1-Phenoxy-2-Propanol in 100 mL of distilled H_2_O (Wyeth et al., 2009). The duration of anaesthesia was determined by testing for a minimal retraction response to manual stimulation of the tentacles and foot. A reduced preparation was dissected while pinned in a Sylgard-lined plastic petri dish (diameter 59 mm) filled with saline (3.00 g/L NaCl, 0.13 g/L KCl, 0.60 g/L CaCl_2_, 0.30 g/L MgCl_2_, 0.61 g/L Tris HCl in dH_2_O, pH 7.4). First, the head and anterior foot were separated from the rest of the body. Then two identical preparations were dissected, each consisting of a half brain, a tentacle nerve, and one tentacle (all still connected within each preparation). For recording, a reduced preparation was transferred to a smaller Sylgard-lined plastic petri dish (diameter 36 mm) and the base of the tentacle was pinned so that the nerve remained accessible to the suction electrode while odour solutions could be delivered to the tentacle. The second preparations from each animal were only ever used for data collection if an inadequate signal-to-noise recording was achieved with the first preparation (and thus only one preparation per animal contributed to our data).

### Recording

Extracellular activity was recorded from the tentacle nerve using a modified *en passant* suction electrode (Johnson et al., 2007). To improve signal to noise ratio via signal averaging, we inserted two silver recording wires (each connected to a separate amplifier) into the barrel of the single polyethylene tubing tip of the electrode. Multiple applications of this twinned electrode to the nerve were typically necessary to obtain optimal signal to noise ratio (assessed qualitatively). The presence of the brain half still attached to the tentacle nerve helped in this regard - it facilitated drawing the longest possible loop of nerve into the polyethylene tip, while plugging the tip opening to create a higher resistance seal. Recordings were obtained using two channels in a A-M Systems Differential AC Amplifier, Model 1700 (settings: 10,000 gain, 10 Hz low-pass filter, 5 kHz high-pass filter), visualized with a Tektronix TDS 204C Four Channel Digital Oscilloscope and digitized at 10 kHz with a Micro1401 analog to digital converter (Cambridge Electronic Design) and Spike2 (version 7) software. In Spike 2, the simultaneous voltage recordings from both electrodes inside the suction tip were averaged. This effectively reduced random noise (which averages towards zero) relative to the electrophysiological signals which were very similar in the two electrodes, differing primarily in the low amplitude (<10 μV) broadband noise, resulting in improved signal-to-noise ratio in comparison to a standard suction electrode with a single recording wire and amplifier (which we had used extensively in pilot work).

### Odour Preparation and Delivery

We tested responses to three odour solutions previously shown to be attractive to *L. stagnalis* (Alansari 2021). The odours were isolated from earthworm-based pellets (Nutrafin Max Bottom Feeder Sinking Food Tablets), algae-based pellets (New Life Spectrum ® AlgaeMax Herbivore Diet 1mm Pellet), and fresh lettuce by soaking the foods in saline. Solutions were made fresh daily and filtered prior to use. The amount of food used to make each solution (22 g earthworm pellet, 22 g algae pellet, 1 cm x 1 cm piece of lettuce) was chosen based on amounts shown in the previous work to stimulate *L. stagnalis* navigational responses (Alansari 2021). Control odour solutions were prepared in identical fashion, but without food in the saline.

Odour solutions were applied to the tentacle preparations using a gravity-fed MPS-2 perfusion system, consisting of a rack with three reservoirs, each connected via polyethylene (PE) tubing to separate cut-off valves and then to a single manifold. The manifold combined fluid flowing from any of the reservoirs into a single stream from a 250 *μ*m blunt-ended needle. The distal end of the needle was positioned in front of the tentacle to ensure a continuous flow of odorants across both the lateral edge and dorsal surface of the organ. The perfusion system design and solenoid operated valves created a negligible time delay when switching between odour solutions. The entire perfusion system was flushed with saline at the beginning and end of each set of trials with a snail. Additionally, each odour treatment solution was always placed in a dedicated reservoir to further avoid any cross contamination between the odour treatments.

### Experimental Design

Three separate experiments were conducted, one for each of the odour sources. Each experiment tested ten snails, with each snail subjected to ten trials (all part of the same recording during a single application of the electrode to the tentacle nerve). The ten trials per recording included five control and five food odour trials presented in random order. Each single trial involved baseline, treatment, and then wash intervals (1 min each). The baseline and wash intervals applied saline to the tentacle, while the treatment interval applied either the control saline or food odour in saline solutions. Baseline intervals were immediately followed by treatment intervals, while wash intervals followed after 15 minutes of constant saline flow (provided from a separate reservoir and tubing system than used with the stimulus manifold) to flush away residual odours (with excess fluid removed by PE tubing connected to a Watson Marlow SciQ 400 403u/vm4 4-channel peristaltic pump). The volume used during flushing was sufficient to completely replace the volume of the recording dish 3 times over.

### Analysis

Using Spike2 software, we averaged the signals from the two silver wires to produce a single waveform for each trial. The 15-minute flushing period between treatment and wash was deleted, and baseline, stimulus and wash intervals were then analyzed similarly. Six voltage thresholds were used to quantify spike activity rates, corresponding to both rising and falling spikes with low, medium, or high amplitudes. Using an algorithm developed from a preliminary experiment (data not shown), thresholds were calculated in an unbiased fashion while allowing for varying signal and noise levels between recordings. The root mean square voltage (RMS) was calculated for all 10 baseline intervals in a recording, and positive and negative thresholds were then calculated at 1.5, 3.3 and 5 times the average baseline RMS level for the recording.

Spike frequencies were calculated for each of the 6 thresholds per interval in all trials. Frequencies were then normalized to the maximum baseline frequency within each set. The normalized data was then used for non-parametric mixed effects factorial analysis with ARTool and corresponding post-hoc comparisons (Wobbrock et al., 2011; Elkin et al. 2021), comparing treatments and intervals while accounting for the lack of independence between intervals and trials conducted on the same preparations. We performed the analyses using Excel (Microsoft Corporation, Redmond, WA, USA), Spike2 (Cambridge Electronic Design, Cambridge, England), and R (Version 4.0.2, R Foundation for Statistical Computing, Vienna, Austria), with R Studio (PBC, Boston, MA, USA) and the following packages: ARTool (Kay et al., 2021), ggplot (Wickham 2016) and ggpubr (Kassambara 2020).

## Results

### Animal-based food odours elicit strong neuronal responses

Odours derived from earthworm-based pellets produced patterns of activity changes that were consistent across all thresholds (Fig. 1,2). Average spike frequencies were similar during the baseline intervals for both control and earthworm-based food odour trials. In control trials, spike frequencies also remained at similar levels during the control odour treatment interval and subsequent wash intervals. In the food odour trials, the treatment interval spike frequencies increased substantially before again reducing frequency close to baseline during the subsequent wash interval. Statistical analysis revealed that, for both low thresholds, most of these differences were significant (Table 1). The only exception was the reduction in frequency during wash intervals following food odour treatment was not significant, presumably because low threshold frequencies were slightly higher during wash intervals than in the baseline intervals.

**Figure 1.**
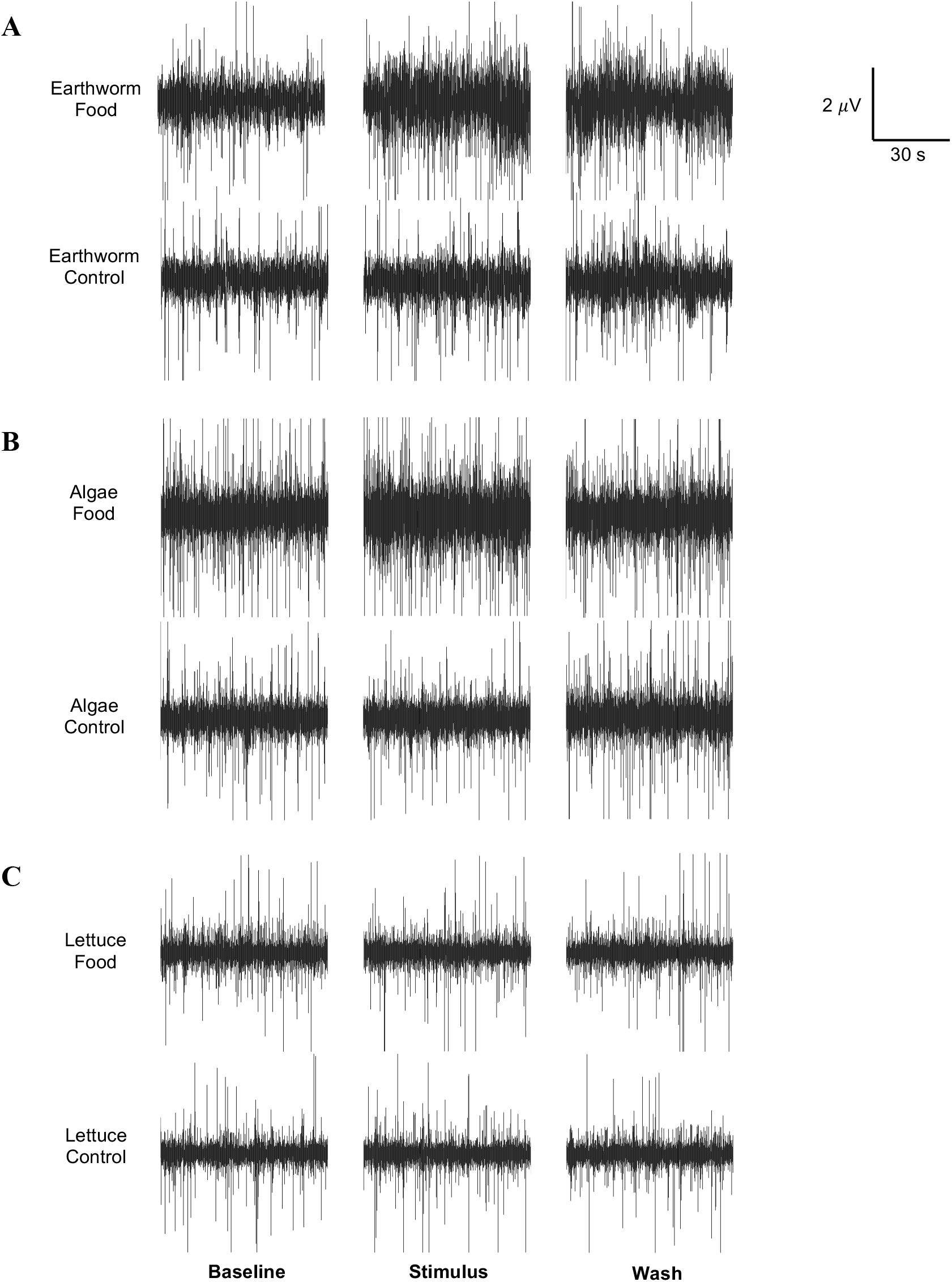
Representative trials from all three experiments show strongest neuronal response to earthworm-based odours. All traces show extracellular responses in the tentacle nerve of *L. stagnalis*. (A) Strong responses were elicited by earthworm odour relative to baseline, wash and control responses. (B) Moderate responses were elicited by algae odour relative to baseline, wash and control responses. (C) Negligible responses were elicited by lettuce odour relative to baseline, wash and control responses.

**Figure 2.**
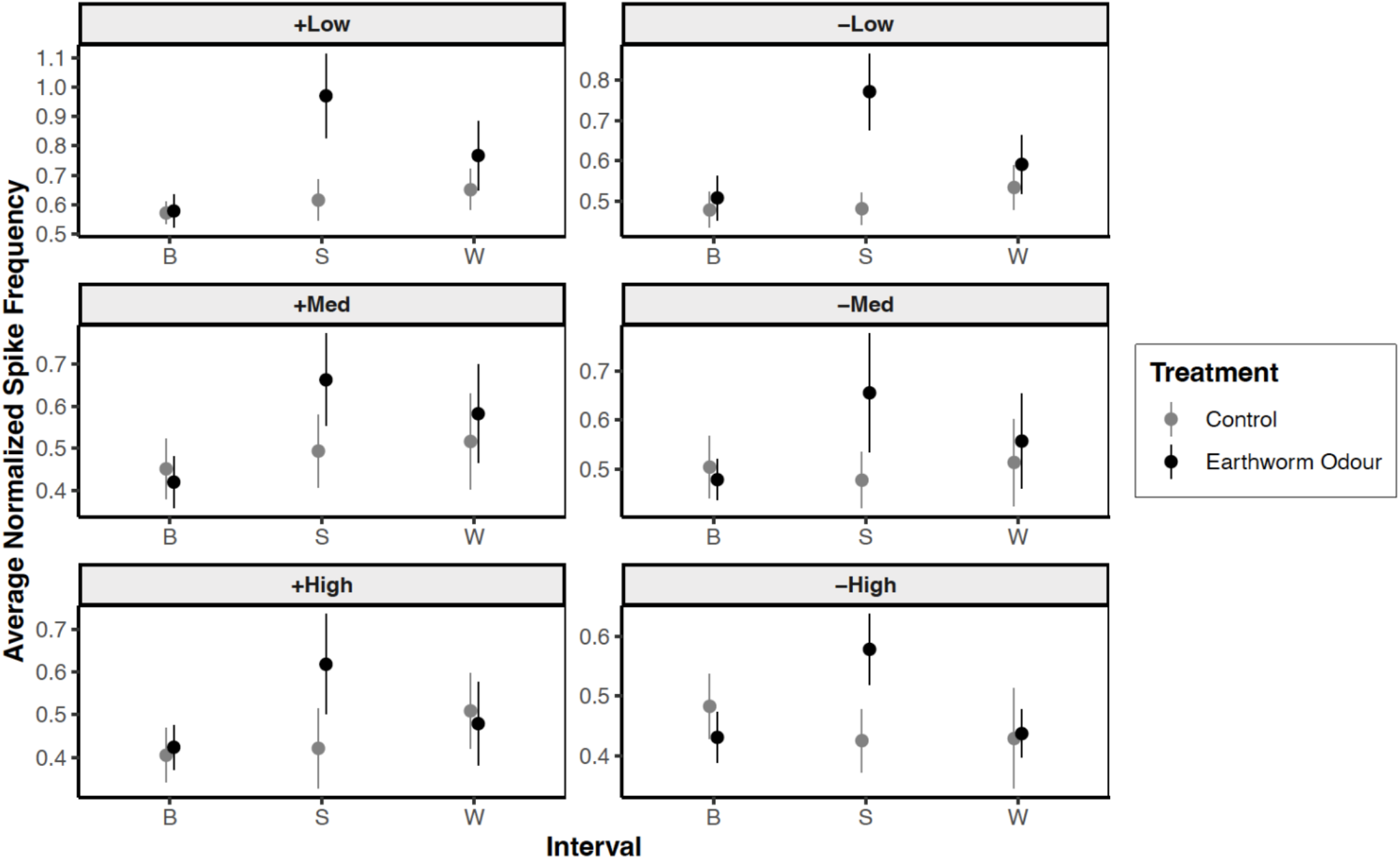
Average spike frequency recorded in the tentacle nerve in response to earthworm-based odours. Each trial included baseline (B), stimulus (S), and wash (W) intervals. Control trials applied saline alone to the tentacle during all three intervals, while food trials applied saline during baseline, the food odour in the stimulus interval, and saline alone again during the wash interval. Each panel presents spike frequencies calculated for one of the six thresholds applied to the voltage waveform data. Each point (+/- s.e.m) indicates the average spike frequency from 10 different preparations.

**Table 1.**
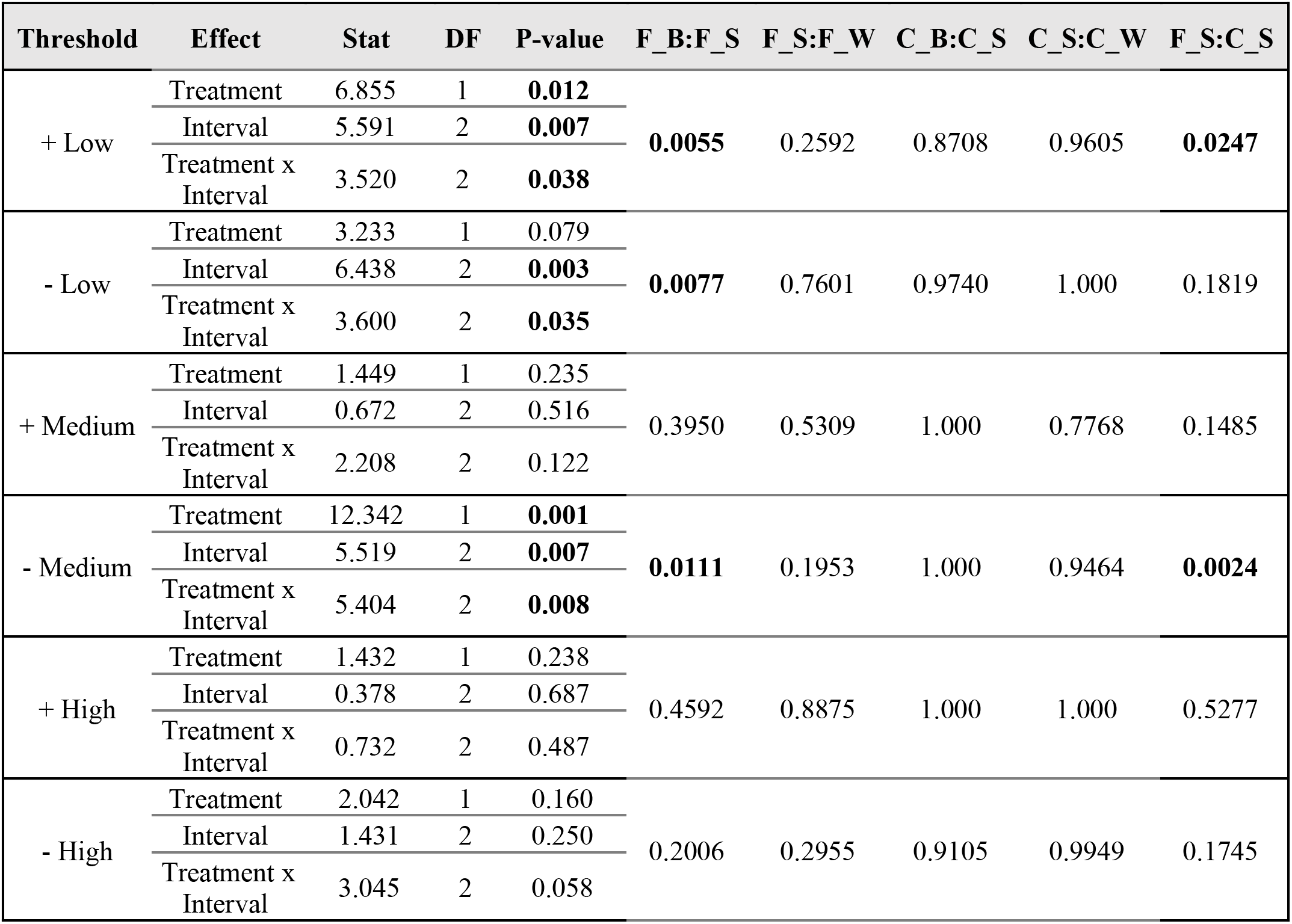
Statistical analysis of spike frequencies across treatments and intervals in response to earthworm-based odours. For each threshold, test statistics, degrees of freedom and *P*-values are reported for the main effects for treatment and interval, and their interaction effect from a two-way mixed effects non-parametric ANOVA. Key post-hoc comparisons between specific treatment interval combinations are reported in the final 5 columns. F_B = food baseline, F_S = food stimulus, F_W =food wash, C_B = control baseline, C_S = control stimulus, C_W = control wash. Bolded numbers indicate *P*-values <0.05.

Although similar patterns occurred with both the medium and high thresholds, only the spike frequencies for the negative medium threshold showed the expected pattern of significant differences between food odour baseline and food odour treatment intervals and between control treatment and food odour treatment intervals. Overall, then, there is clear evidence for a neuronal response to earthworm odour quantified by the low thresholds. The medium and high thresholds have weaker evidence, although the expected trends in spike frequencies were all still present.

### Algae-based food odours elicit weak neuronal responses

In this experiment, the patterns of activity changes between intervals and treatments were again consistent with expectations if the algae-based food odours stimulated chemosensory activity in the tentacle nerve. However, only some of the expected differences in activity were significant, and only for the low threshold data (Fig. 3). Average spike frequencies were similar in the baseline interval of both the control and algae-based food odour treatment with all thresholds. In the control treatment, there was a similar average spike frequency across the stimulus and wash intervals. In the food treatment, the average spike frequency increased as algae-based odours were applied during the stimulus interval, followed by a subsequent decrease in average spike frequency during the wash interval. However, clear trends suggesting a return to baseline frequencies were only present in the positive threshold data. Statistical analysis only revealed significant changes from baseline during the food stimulus interval, and only with the low thresholds for analysis (Table 2). Otherwise, no significant differences in spike frequency occurred either between control and food treatment stimulus intervals (despite the strong trends in the low threshold data), nor between stimulus and wash intervals. These data are all consistent with a neuronal response to algae-based odours, with spike amplitudes and frequencies such that they are not as distinct from the background noise as with the animal-food based odours.

**Figure 3.**
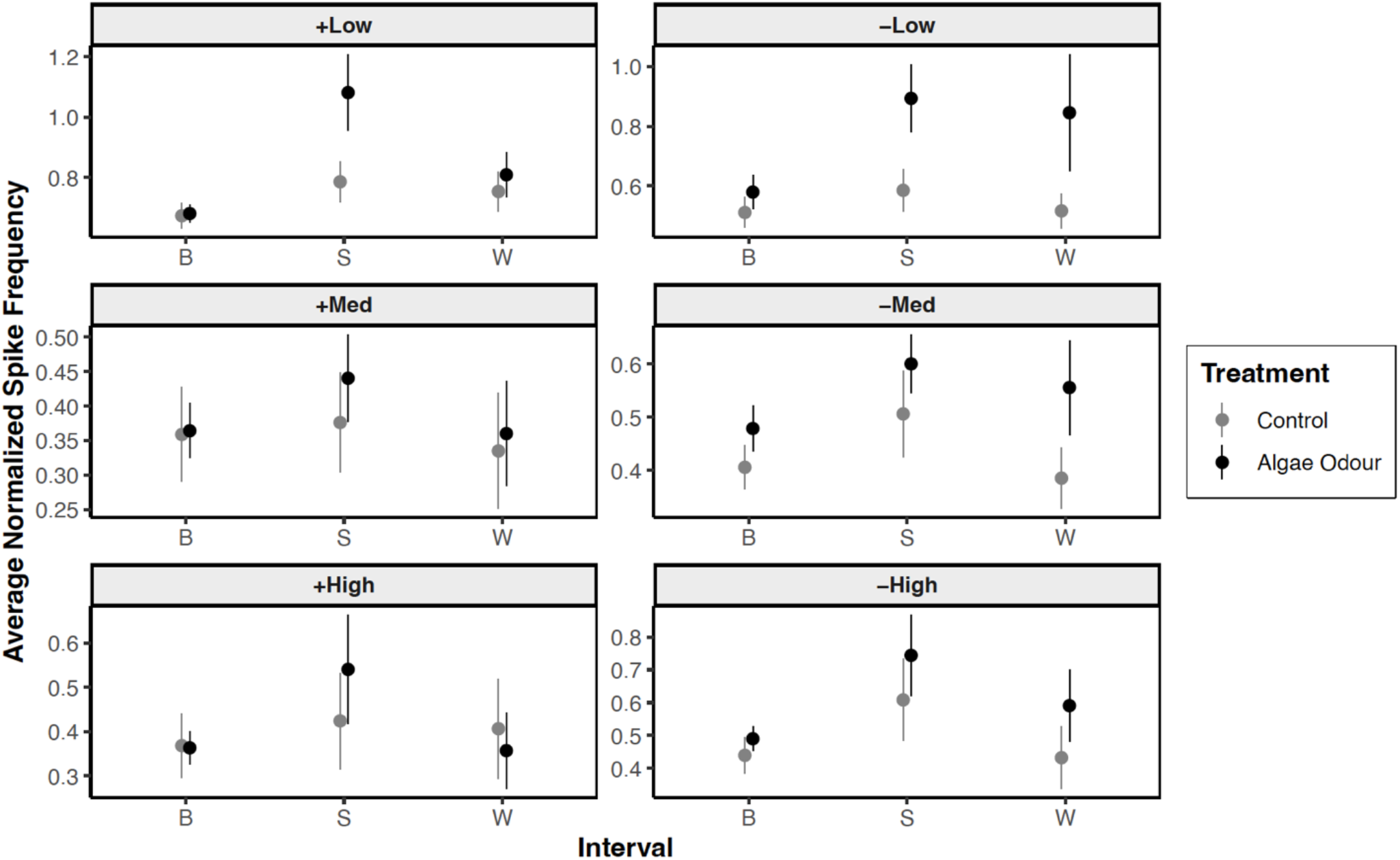
Average spike frequency recorded in the tentacle nerve in response to algae-based odours. Each trial included baseline (B), stimulus (S), and wash (W) intervals. Control trials applied saline alone to the tentacle during all three intervals, while food trials applied saline during baseline, the food odour in the stimulus interval, and saline alone again during the wash interval. Each panel presents spike frequencies calculated for one of the six thresholds applied to the voltage waveform data. Each point (+/- s.e.m) indicates the average spike frequency from 10 different preparations.

**Figure 4.**
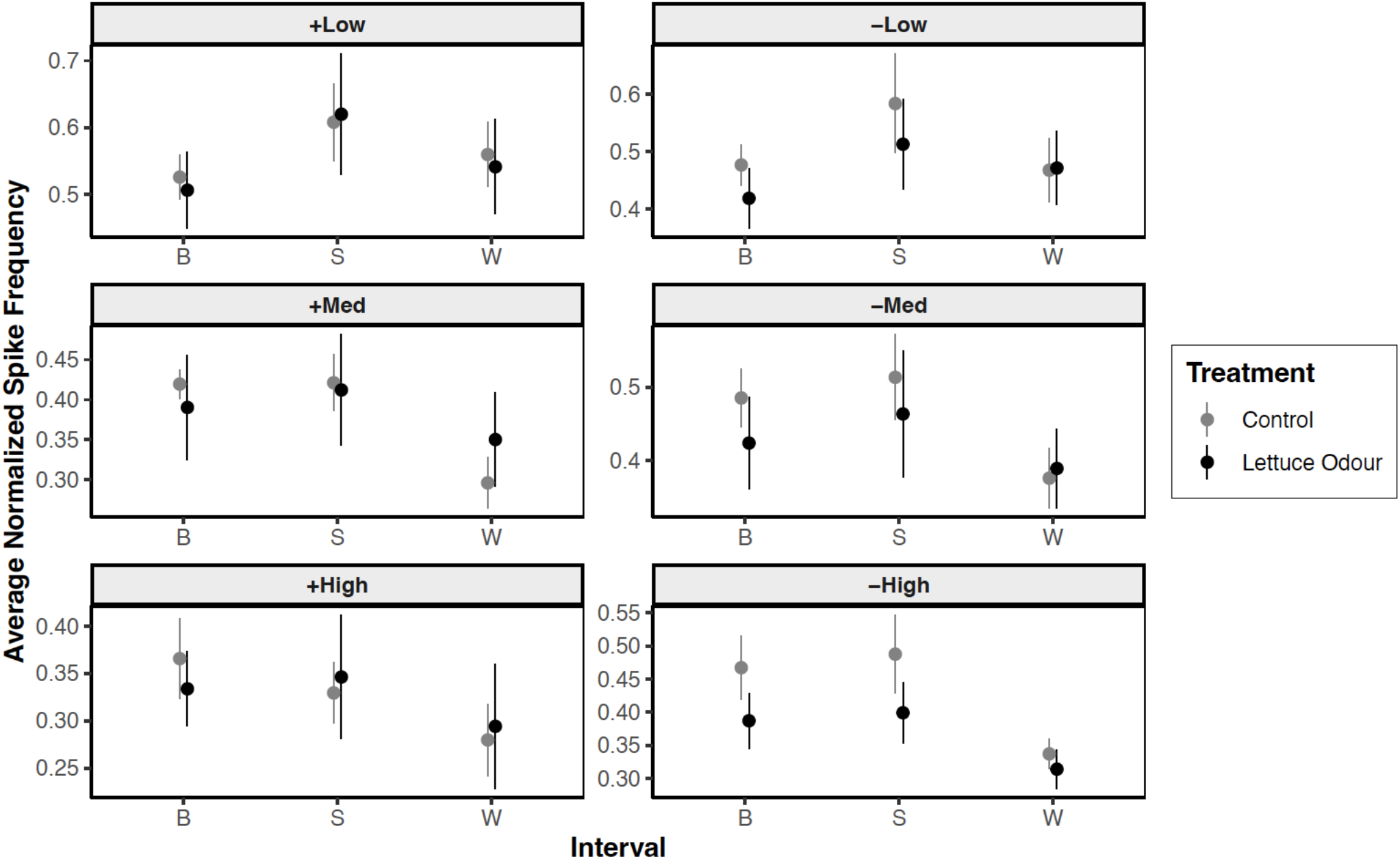
Average spike frequency recorded in the tentacle nerve in response to lettuce odours. Each trial included baseline (B), stimulus (S), and wash (W) intervals. Control trials applied saline alone to the tentacle during all three intervals, while food trials applied saline during baseline, the food odour in the stimulus interval, and saline alone again during the wash interval. Each panel presents spike frequencies calculated for one of the six thresholds applied to the voltage waveform data. Each point (+/- s.e.m) indicates the average spike frequency from 10 different preparations.

**Table 2.** Statistical analysis of spike frequencies across treatments and intervals in response to algae-based odours. For each threshold, test statistics, degrees of freedom and *P-*values are reported for the main effects for treatment and interval, and their interaction effect from a two-way mixed effects non-parametric ANOVA. Key post-hoc comparisons between specific treatment interval combinations are reported in the final 5 columns. F_B = food baseline, F_S = food stimulus, F_W =food wash, C_B = control baseline, C_S = control stimulus, C_W = control wash. Bolded numbers indicate *P*-values <0.05.

| Threshold | Effect | Stat | DF | P-value | F_B:F_S | F_S:F_W | C_B:C_S | C_S:C_W | F_S:C_S |
| --- | --- | --- | --- | --- | --- | --- | --- | --- | --- |
| + Low | Treatment | 12.556 | 1 | **<0.001** | **<0.001** | 0.142 | 0.060 | 0.851 | 0.179 |
|  | Interval | 20.418 | 2 | **<0.001** |  |  |  |  |  |
|  | Treatment x Interval | 5.939 | 2 | **0.005** |  |  |  |  |  |
| - Low | Treatment | 15.228 | 1 | **<0.001** | **0.0397** | 0.692 | 0.846 | 0.973 | 0.110 |
|  | Interval | 6.652 | 2 | **0.003** |  |  |  |  |  |
|  | Treatment x Interval | 3.083 | 2 | 0.056 |  |  |  |  |  |
| + Medium | Treatment | 1.364 | 1 | 0.249 | 0.723 | 0.708 | 1.000 | 0.947 | 0.806 |
|  | Interval | 1.737 | 2 | 0.188 |  |  |  |  |  |
|  | Treatment x Interval | 0.252 | 2 | 0.778 |  |  |  |  |  |
| - Medium | Treatment | 11.402 | 1 | **0.002** | 0.450 | 0.934 | 0.726 | 0.583 | 0.393 |
|  | Interval | 2.163 | 2 | 0.127 |  |  |  |  |  |
|  | Treatment x Interval | 0.705 | 2 | 0.500 |  |  |  |  |  |
| + High | Treatment | 0.829 | 1 | 0.367 | 0.765 | 0.352 | 1.000 | 1.000 | 0.628 |
|  | Interval | 1.424 | 2 | 0.251 |  |  |  |  |  |
|  | Treatment x Interval | 0.883 | 2 | 0.421 |  |  |  |  |  |
| - High | Treatment | 5.631 | 1 | **0.022** | 0.349 | 0.703 | 0.662 | 0.386 | 0.689 |
|  | Interval | 3.622 | 2 | **0.035** |  |  |  |  |  |
|  | Treatment x Interval | 0.396 | 2 | 0.675 |  |  |  |  |  |

**Table 3.** Statistical analysis of spike frequencies across treatments and intervals in response to lettuce odours. For each threshold, test statistics, degrees of freedom and *P-*values are reported for the main effects for treatment and interval, and their interaction effect from a two-way mixed effects non-parametric ANOVA. Key post-hoc comparisons between specific treatment interval combinations are reported in the final 5 columns. F_B = food baseline, F_S = food stimulus, F_W =food wash, C_B = control baseline, C_S = control stimulus, C_W = control wash. Bolded numbers indicate *P*-values <0.05.

| Threshold | Effect | Stat | DF | P-value | F_B:F_S | F_S:F_W | C_B:C_S | C_S:C_W | F_S:C_S |
| --- | --- | --- | --- | --- | --- | --- | --- | --- | --- |
| + Low | Treatment | 0.228 | 1 | 0.635 | 0.805 | 0.992 | 0.603 | 0.991 | 0.999 |
|  | Interval | 2.133 | 2 | 0.130 |  |  |  |  |  |
|  | Treatment x Interval | 0.037 | 2 | 0.964 |  |  |  |  |  |
| - Low | Treatment | 0.750 | 1 | 0.391 | 0.888 | 1.000 | 0.619 | 0.718 | 0.903 |
|  | Interval | 1.847 | 2 | 0.170 |  |  |  |  |  |
|  | Treatment x Interval | 0.337 | 2 | 0.715 |  |  |  |  |  |
| + Medium | Treatment | 0.290 | 1 | 0.593 | 0.991 | 0.904 | 1.000 | **0.019** | 0.930 |
|  | Interval | 4.261 | 2 | **0.020** |  |  |  |  |  |
|  | Treatment x Interval | 1.243 | 2 | 0.298 |  |  |  |  |  |
| - Medium | Treatment | 1.127 | 1 | 0.294 | 0.996 | 0.805 | 0.999 | **0.046** | 0.927 |
|  | Interval | 3.867 | 2 | **0.028** |  |  |  |  |  |
|  | Treatment x Interval | 0.684 | 2 | 0.510 |  |  |  |  |  |
| + High | Treatment | 0.294 | 1 | 0.590 | 0.999 | 0.969 | 0.990 | 0.906 | 1.000 |
|  | Interval | 1.694 | 2 | 0.195 |  |  |  |  |  |
|  | Treatment x Interval | 0.014 | 2 | 0.986 |  |  |  |  |  |
| - High | Treatment | 6.733 | 1 | **0.013** | 1.000 | 0.333 | 1.000 | 0.110 | 0.667 |
|  | Interval | 7.927 | 2 | **0.001** |  |  |  |  |  |
|  | Treatment x Interval | 0.767 | 2 | 0.471 |  |  |  |  |  |

**Table 4.**
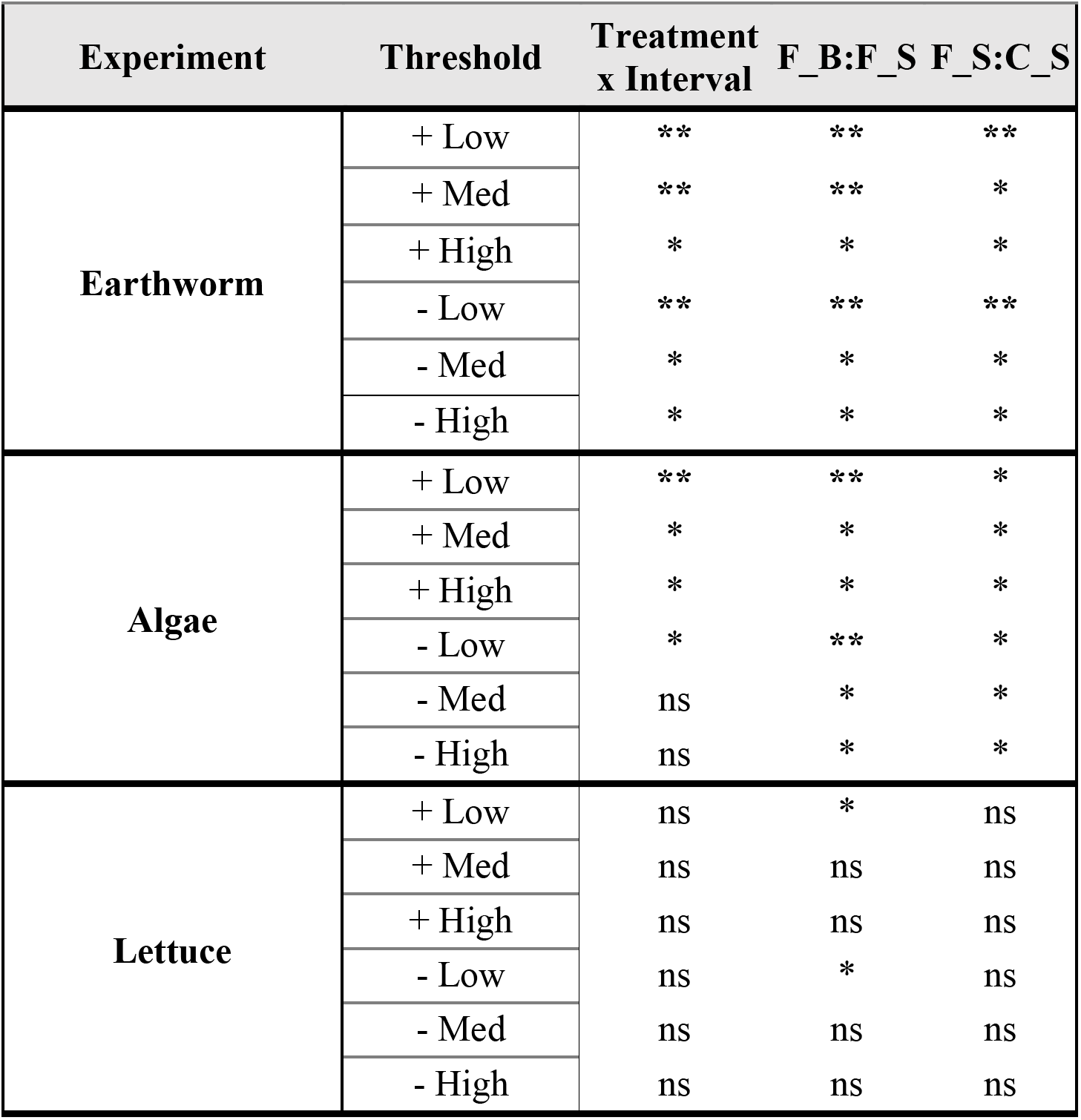
Summary of the major results from all three odour experiments 595 for all thresholds analyzed. The key statistical results supporting the hypothesis that the odour solution produced a chemosensory response were: 1) a significant interaction effect (Treatment x Interval), with post-hoc comparisons showing a difference between food baseline and food stimulus (F_B:F_S) as well as a difference between food stimulus and control stimulus (F_S:C_S). (**) indicates statistically significant trends consistent with a chemosensory response to the food odour, (*) indicates a non-significant trend, and (ns) represents no such trend.

### Lettuce odours do not elicit neuronal responses

## Discussion

We recorded extracellular responses in the tentacle nerves of *L. stagnalis* to two of the three different food odours we tested. The strongest responses were elicited by odours derived from earthworm-based fish food pellets. Spike frequencies recorded in the tentacle nerve increased only when the odours were applied (and not during control applications without odours) and diminished again after wash (Fig. 1). These differences in spike frequency were clearest with relatively low thresholds applied to the recordings but were also evident when a medium threshold was applied. This suggests that many responding neurons produced relatively low amplitude spikes, presumably originating in a population of neurons with small diameter axons. Algae-based food odours stimulated similar, albeit weaker neural responses. Finally, we did not record any significant neural response to lettuce odours in the tentacle nerve. It is not yet clear whether these differences in responses among the odours we tested were a consequence of differing sizes, numbers, or populations of responding neurons.

The differing responses to the three odours used in this study may be caused by several reasons. (1) We cannot eliminate the possibility that different response strengths occurred because of variations in the concentration of odour molecules present in each stimulus (Gillette et al. 2000; Noboa and Gillette 2013). Since both pellet and fresh foods were used to generate the odours, there are likely differences in the concentrations, the release rates, and the composition of odorants present in each odour stimulus. It is possible that the artificial food sources produced a higher concentration of stimulating odorants in solution, leading to enhanced neuronal responses in the earthworm and algae treatments compared to the lettuce treatment. (2) The design of our study also provided no insight into the number of different chemosensory neurons in the tentacles that may be responding to odours. Along with variations in the number of chemosensory neurons, it is possible that different odours or different combinations of odours in each food treatment stimulate a separate set of chemosensory neurons. Thus, both the number and type of chemosensory neurons present in the tentacles may have influenced the neuronal activity generated during the olfactory responses. (3) If different odours generate neuronal signals with varying amplitudes and frequencies, there is a potential for varying levels of signal exclusion from our recordings. Thus, for example, if the signals associated with lettuce odour stimuli were sufficiently smaller than the signals elicited by the other food odours, this could explain the absence of any responses during the lettuce experiment. Indeed, our pilot work with a traditional single channel suction electrode found only partial evidence of responses to earthworm odours, and it was only improving signal-to-noise ratio with our twinned electrode that the results presented here were obtained. Consequently, we cannot definitively conclude that *L. stagnalis* does not respond to lettuce odours that would be revealed by further improvements to the recording system.

The relative responses to different food odour types recorded here are consistent with previous work assessing foraging responses in *L. stagnalis*. Early studies of *L. stagnalis* report its omnivorous feeding habits. This grazing gastropod feeds primarily on freshwater aquatic flora, such as *Elodea sp*. and algae biofilms, but will also feed on carrion when it is available (Balance et al. 2001; Bovberg 1968). Behavioral studies suggest a preference for carrion, as *L. stagnalis* abandoned feeding on plant food for carrion and showed the strongest navigational responses to the same earthworm-based food pellets used here (Bovberg 1968; Alansari 2021). The results of our study are consistent with this behavioral finding, showing a stronger neuronal response to earthworm pellet odours than algae pellet and lettuce odours. It may be higher fitness for *L. stagnalis* to respond more strongly to animal-based odours. Animal carrion can be a highly nutritious but rare food source compared to plants (Parmenter and MacMahon 2009). Given the high reward of feeding on animal carrion, it may be beneficial for the snails to seek out this food source. Additionally, since plants offer less nutrition and are abundant in the environment, *L. stagnalis* may not employ olfactory driven navigational strategies in search of plant food sources. Instead, the snail may resort to feeding on plants in the absence of animal carrion, perhaps during (or after) failed foraging attempts on animal carrion. Given that lettuce is not a natural food source available in the habitat of *L. stagnalis*, it is also possible that these animals do not have sensory receptors able to respond to lettuce odours. While a study has reported that lettuce is a good alternative food source (Calow 1973) and the plant is used as a primary diet for snails in laboratories, this utility may not necessarily correlate with the appropriate adaptation for navigation towards the food source. Regardless, our data provides evidence that *L. stagnalis* responds more strongly to animal food sources than to plant-based odours.

Our extracellular recordings from the tentacle nerve of *L. stagnalis* confirmed that the tentacles are chemosensory. Both the tentacles and lips comprise the cephalic sensory organs, which are typically considered to bear chemosensory and mechanosensory neurons in gastropods (Cummins and Wyeth 2014). In *L. stagnalis*, neuroanatomical studies showed an extensive distribution of peripheral sensory neurons across the tentacles (and the lips). (Wyeth and Croll 2011; Horvath et al. 2020). Previous electrophysiological studies in *L. stagnalis* have shown neuronal responses to both tactile and sugar stimuli of the lips or buccal mass (Jager 1971; Goldschmeding and Jager 1973; Kemenes et al. 1986; Nakamura et al. 1999). Although responses to KCl could be recorded from the tentacle nerve, tentacle responses to sugar stimuli were absent (Nakamura et al. 1999). These results are consistent with sugar acting as a gustatory stimulus leading to feeding responses via sensory receptors present only in the lips. Our study used only the tentacles and a true olfactory stimulus (solely odours carried by the fluid medium from a distant odour source). We were able to record clear differences in neuronal signalling in response to two of the three odours, indicating the chemosensory nature of the tentacles, and further suggesting their probable role in olfactory-based navigation relative to these and presumably other odours.

Our results suggest peripheral sensory cells may be the first component of the olfactory system to distinguish between odour stimuli. Electrophysiological studies in vertebrates and invertebrates using odor mixtures and their components have revealed potential peripheral decoding of sensory signals (Thomas-Danguin et al. 2014). Still, the process of odour differentiation, particularly in the context of complex stimuli that contain more than one odorant, is not well understood. As our study shows that afferent activity in the tentacle nerve responds most strongly to earthworm-based food odours, this would suggest that early odour differentiation may begin at the peripheral nervous system level with chemosensory neurons in the tentacle. However, it is critical that we continue to consider whether the differing neuronal responses are because of the odor concentration or the type and number of chemosensory neurons in the cephalic sensory organs. The snail may differentiate between odours based solely on the concentrations of odorants present in all stimuli, resulting in solely firing rate differences in the response to different odours. On the other hand, odour differentiation may rely on the type or number of chemosensory neurons that are being stimulated at any given time by differing mixtures of odorants. Both options may be potential mechanisms for peripheral odour differentiation in *L. stagnalis*. Although it is not yet clear what contributed to the different neuronal responses in this study, our results nonetheless point toward early odour differentiation beginning in the peripheral nervous system of the tentacles.

There remain several further unanswered questions regarding the mechanisms by which *L. stagnalis* detects, integrates, and responds to odour cues. Are the lips involved in distance chemoreception in conjunction with the tentacles? Similar experiments as used here but involving the lips will provide insight into the chemosensory and possible olfactory function of this cephalic sensory organ. Further work is also needed to identify the nature and concentrations of odorants released by the different odour sources tested here, and which of those molecules are stimulating tentacular chemosensory neurons. Nonetheless, our study adds our understanding of chemosensation in gastropods and provides the foundation for future research into the neuroethology of odour-based navigation in *L. stagnalis*. Finally, we also suggest our twinned electrode recording system followed by signal averaging may be useful for extracellular recordings form other preparations with small amplitude signals.

## Acknowledgments

We thank Areej Alansari for support during the initial stages of this study, and the staff of the Animal Care Facility at St. Francis Xavier University.

